# Consortium genome-wide meta-analysis for childhood dental caries traits

**DOI:** 10.1101/238824

**Authors:** Simon Haworth, Dmitry Shungin, Justin T van der Tas, Strahinja Vucic, Carolina Medina Gomez, Victor Yakimov, Bjarke Feenstra, John R Shaffer, Myoung Keun Lee, Marie Standl, Elisabeth Thiering, Carol Wang, Klaus Bønnelykke, Johannes Waage, Leon Eyrich Jessen, Pia Elisabeth Nørrisgaard, Raimo Joro, Ilkka Seppälä, Olli Raitakari, Tom Dudding, Olja Grgic, Edwin Ongkosuwito, Anu Vierola, Aino-Maija Eloranta, Nicola X West, Steven J Thomas, Daniel W McNeil, Steven M Levy, Rebecca Slayton, Ellen A Nohr, Terho Lehtimäki, Timo Lakka, Hans Bisgaard, Craig Pennell, Jan Kühnisch, Mary L Marazita, Mads Melbye, Frank Geller, Fernando Rivadeneira, Eppo B Wolvius, Paul W Franks, Ingegerd Johansson, Nicholas J Timpson

## Abstract

Prior studies suggest dental caries traits in children and adolescents are partially heritable, but there has been no large-scale consortium genome-wide association study (GWAS) to date. We therefore performed GWAS for caries in participants aged 2.5-18.0 years from 9 contributing centers. Phenotype definitions were created for the presence or absence of treated or untreated caries, stratified by primary and permanent dentition. All studies tested for association between caries and genotype dosage (imputed to Haplotype Reference Consortium or 1000 Genomes phase 1 version 3 panels) accounting for population stratification. Fixed–effects meta-analysis was performed weighted by inverse standard error. Analysis included up to 19,003 individuals (7,530 affected) for primary teeth and 13,353 individuals (5,875 affected) for permanent teeth. Evidence for association with caries status was observed at rs1594318-C for primary teeth (intronic within *ALLC*, Odds Ratio (OR) 0.85, Effect Allele Frequency (EAF) 0.60, p 4.13e-8) and rs7738851-A (intronic within *NEDD9*, OR 1.28, EAF 0.85, p 1.63e-8) for permanent teeth. Consortium-wide estimated heritability of caries was low (h^2^ of 1% [95% CI: 0%:7%] and 6% [95% CI 0%:13%] for primary and permanent dentitions, respectively) compared to corresponding within-study estimates (h^2^ of 28%, [95% CI: 9%:48%] and 17% [95% CI:2%:31%]) or previously published estimates. This study was designed to identify common genetic variants with modest effects which are consistent across different populations. We found few single variants associated with caries status under these assumptions. Phenotypic heterogeneity between cohorts and limited statistical power will have contributed; these findings could also reflect complexity not captured by our study design, such as genetic effects which are conditional on environmental exposure.

**Author summary:** Dental caries (tooth decay) is a common disease in children. Previous studies suggest genetic factors alter caries risk, but to date there is a gap of knowledge in identifying which specific genetic variants are responsible. We undertook analysis in a consortium including around 19,000 children and investigated whether any of 8 million common genetic variants were associated with risk of caries in primary (milk) or permanent teeth. If identified, these variants are used as ‘tags’ to highlight genes which may be involved in a disease. We identified variants in two loci associated with caries status; in the primary (rs1594318) and permanent dentition (rs7738851). The former is intronic in *ALLC*, a gene with poorly understood function. The latter is an intronic variant within *NEDD9*, a gene which has several known functions including a role in development of craniofacial structures. To gain a more comprehensive understanding of genetic effects which influence caries larger studies and a better understanding of environmental modifiers or interactions with genetic effects are required.

## Introduction

Dental caries remains a prevalent public health problem in both children and adults. Untreated dental caries was estimated to affect 621 million children worldwide in 2010, with little change in prevalence or incidence between 1990 and 2010 [1]. This problem is not unique to lower income countries; around 50% of children have evidence of caries by age 5 in industrialized nations [2–4]. Dental caries results from reduced mineral saturation of fluids surrounding teeth, driven by acid produced by bacteria in the oral microbiome [5]. Many different factors predispose towards dental caries, of which high sugar consumption, poor oral hygiene and low socio-economic status are the most notorious [5–8]. Over the last decades there has been increasing appreciation for the role of genetic influences in dental caries. The importance of genetic susceptibility for dental caries experience was demonstrated in an animal model over 50 years ago, a finding since substantiated in twin studies in humans [9–11]. Of particular relevance to caries traits in children and adolescents, Bretz et al. analyzed longitudinal rates of change in caries status in children, and found that caries progression and severity were highly heritable in the primary and permanent dentition [10]. It has also been suggested that heritability for dental caries does not depend entirely on genetic predisposition to sweet food consumption [12]. Despite evidence of a genetic contribution to caries susceptibility, few specific genetic loci have been identified.

Shaffer et al. performed the first GWAS for dental caries in 2011 [13], studying the primary dentition of 1,305 children. They found evidence for association at novel and previously studied candidate genes *(ACTN2, MTR, EDARADD, MPPED2* and *LPO)*, but no individual single nucleotide polymorphisms (SNPs) exceeded the genome wide significance threshold (P-value ≤ 5.0E-08), possibly as a consequence of the modest sample size [13]. The first GWAS for dental caries in the permanent dentition in adults was performed at a similar time by Wang et al [14]. They included 7,443 adults from five different cohorts and identified several suggestive loci (P-value ≤ 10E-05) for dental caries *(RPS6KA2, PTK2B, RHOU, FZD1, ADMTS3* and *ISL1*), different loci from those mentioned above for the primary dentition and again with no single variants reaching genome-wide significance.

The next wave of GWAS of caries suggested association at a range of different loci. Two GWAS used separate phenotype definitions for pit-and-fissure and smooth tooth surfaces and identified different loci associated with dental caries susceptibility in both primary and permanent dentition [15, 16]. The GWAS in primary dentition used a sample of approximately 1,000 children and found evidence for association at loci reported in previous studies, including *MPPED2, RPS6KA2*, and *AJAP1* [13–16]. The largest GWAS for dental caries in permanent dentition was performed in a Hispanic and Latino sample of 11,754 adults [17]. This study identified unique genetic loci (*NAMPT* and *BMP7*) compared to previous GWAS in individuals of European ancestry. To date it is unclear whether the variability in nominated loci reflects true variability in the genetic architecture of dental caries across different populations, age periods and sub-phenotypic definitions, or merely represent chance differences between studies given the modest power in the studies performed to date.

Dental caries is a complex and multifactorial disease, caused by a complex interplay between environmental, behavioral and genetic factors. Until now there has been a lack of large scale studies of dental caries traits in children and the genetic basis of these traits remains poorly characterized. This investigation set out to examine the hypothesis that common genetic variants influence dental caries with modest effects on susceptibility. We anticipated that a) caries in both primary and permanent teeth would be heritable in children and adolescents aged 2.5 – to - 18 years and b) common genetic variants are likely to only have small effects on the susceptibility of a complex disease such as dental caries. Therefore, the aim of this large-scale, consortium-based GWAS is to examine novel genetic loci associated with dental caries in primary and permanent dentition in children and adolescents.

## Methods

### Study samples

We performed genome-wide association (GWA) analysis for dental caries case/control status in a consortium including 9 coordinating centers. Study procedures differed between these centers. We use the term ‘clinical dental assessment’ to mean that a child was examined in person, whether this was in a dental clinic or a study center. We use the term ‘examiner’ to refer to a dental professional, and use the term ‘assessor’ to refer to an individual with training who is not a dental professional, for example a trained research nurse.

The Avon Longitudinal Study of Parents and Children (ALSPAC) is a longitudinal birth cohort which recruited pregnant women living near Bristol, UK with an estimated delivery date between 1991 and 1992. Follow up has included clinical assessment and questionnaires and is ongoing [18]. A subset of children attended clinics including clinical dental assessment by a trained assessor at age 31, 43 and 61 months of age. Parents were asked to complete questionnaires about their children’s health regularly, including comprehensive questions at a mean age of 5.4 and 6.4 years. Parents and children were asked to complete questionnaires about oral health at a mean age of age 7.5, 10.7 and 17.8 years. Please note that the study website contains details of all the data that are available through a fully searchable data dictionary (http://www.bris.ac.uk/alspac/researchers/data-access/data-dictionary/). Both clinical and questionnaire derived data were included in this analysis, with priority given to clinical data where available. (S3 Table).

The Copenhagen Prospective Studies on Asthma in Childhood includes two population based longitudinal birth cohorts in Eastern Denmark. C0PSAC2000 recruited pregnant women with a history of asthma between 1998 and 2001 [19]. Children who developed wheeze in early life were considered for enrollment in a nested randomized trial for asthma prevention. COPSAC2010 recruited pregnant women between 2008 and 2010 and was not selected on asthma status. Both COPSAC2000 and COPSAC2010 studies included regular clinical follow up. Within Denmark clinical dental assessment is routinely offered to children and adolescents until the age of 18 years and summary data from these examinations are stored in a national register. These data were obtained via index linkage for participants of COPSAC2000 and COPSAC2010 and used to perform joint analysis across both cohorts.

The Danish National Birth Cohort (DNBC) is a longitudinal birth cohort which recruited women in midpregnancy from 1996 onwards [20]. For this analysis, index linkage was performed to obtain childhood dental records for mothers participating in DNBC. As with the COPSAC studies, these data were originally obtained by a qualified dentist and included surface level dental charting.

The Generation R study (GENR) recruited women in early pregnancy with expected delivery dates between 2002 and 2006 living in the city of Rotterdam, the Netherlands. The cohort is multi-ethnic with representation from several non-European ethnic groups. Follow-up has included clinical assessment visits and questionnaires and is ongoing [21, 22]. Intra-oral photography was performed as a part of their study protocol, with surface level charting produced by a dental examiner (a specialist in pediatric dentistry) [23]. Analysis in GENR included a) a multi-ethnic association study including all individuals with genetic and phenotypic data and b) analysis including only individuals of European ancestry.

The GENEVA consortium is a group of studies which undertake coordinated analysis across several phenotypes. [24] Within GENEVA, the Center for Oral Health Research in rural Appalachia, West Virginia and Pennsylvania, USA (COHRA), the Iowa Fluoride Study in Iowa, USA (IFS) and the Iowa Head Start (IHS) study participated in analysis of dental traits in children [25]. COHRA recruited families with at least one child aged between 1 and 18 years of age, with dental examination performed at baseline [26]. IFS recruited mothers and newborn infants in Iowa between 1992 and 1995 with a focus on longitudinal fluoride exposures and dental and bone health outcomes. Clinical dental examination in IFS was performed by trained assessors age 5, 9 13 and 17 years [27]. IHS recruited children participating in an early childhood education program which included a one-time clinical dental examination [13].

The “German Infant study on the influence of Nutrition Intervention plus air pollution and genetics on allergy development” (GINIplus) is a multi-center prospective birth cohort study which has an observational and interventional arm which conducted a nutritional intervention during the first four months of life. The study recruited new born infants with and without family history of allergy in the Munich and Wesel areas, Germany between 1995 and 1998 [28, 29]. The “Lifestyle-related factors, Immune System and the development of Allergies in East and West Germany” study (LISA) is a longitudinal birth cohort which recruited between 1997 and 1999 across four sites in Germany [28, 30]. For participants living in the Munich area, follow up used similar protocols in both GINIplus and LISA, with questionnaire and clinic data including clinical dental examination by trained examiners at age 10 and 15 years. Analysis for caries in GINIplus and LISA was therefore performed across both studies for participants at the Munich study center.

The Physical Activity and Nutrition in Children (PANIC) Study is an ongoing controlled physical activity and dietary intervention study in a population of children followed retrospectively since pregnancy and prospectively until adolescence. Altogether 512 children 6-8 years of age were recruited in 2008-2009 [31]. The main aims of the study are to investigate risk factors and pathophysiological mechanisms for overweight, type 2 diabetes, atherosclerotic cardiovascular diseases, musculoskeletal diseases, psychiatric disorders, dementia and oral health problems and the effects of a long-term physical activity and dietary intervention on these risk factors and pathophysiological mechanisms. Clinical dental examinations were performed by a qualified dentist with tooth level charting.

The Cardiovascular Risk in Young Finns Study (YFS) is a multi-center investigation which aimed to understand the determinants of cardiovascular risk factors in young people in Finland. The study recruited participants who were aged 3, 6, 9, 12, 15 and 18 years old in 1980. Eligible participants living in specific regions of Finland were identified at random from a national population register and were invited to participate. Regular follow-up has been performed through physical examination and questionnaires [32]. Clinical dental examination was performed by a qualified dentist with tooth level charting.

The Western Australian Pregnancy Cohort (RAINE) study is a birth cohort which recruited women between 16^th^ and 20^th^ week of pregnancy living in the Perth area, Western Australia. Recruitment occurred between 1989 and 1991 with regular follow up of mothers and their children through research clinics and questionnaires [33]. The presence or absence of dental caries was recorded by a trained assessor following clinical dental examination at the year 3 clinic follow up.

Further details of study samples are provided in S1 Table.

### Medical Ethics

Within each participating study written informed consent was obtained from the parents of participating children after receiving a full explanation of the study. Children were invited to give assent where appropriate. All studies were conducted in accordance with the Declaration of Helsinki.

Ethical approval for the ALSPAC study was obtained from the ALSPAC Ethics and Law Committee and the Local Research Ethics Committee. Full details of ethical approval policies and supporting documentation are available online (http://www.bristol.ac.uk/alspac/researchers/research-ethics/.) Approval to undertake analysis of caries traits was granted by the ALSPAC executive committee (B2356).

The COPSAC2000 cohort was approved by the Regional Scientific Ethical Committee for Copenhagen and Frederiksberg (KF 01-289/96) and the Danish Data Protection Agency (2008-41-1574). The 2010 cohort (COPSAC2010) was approved by the Danish Ethics Committee (H-B-2008-093) and the Danish Data Protection Agency (2008-41-2599).

The DNBC study of caries was approved by the Scientific Ethics Committee for the Capital City Region (Copenhagen), the Danish Data Protection Agency, and the DNBC steering committee.

Each participating site in the GENEVA consortium caries analysis received approval from the local university institutional review board (federal wide assurance number for GENEVA caries project: FWA00006790). Within the COHRA arm local approval was provided by the University of Pittsburgh (020703/0506048) and West Virginia University (15620B), whilst the IFS and IHS arms received local approval from the University of Iowa’s Institutional Review Board.

The GENR study design and specific data acquisition were approved by the Medical Ethical Committee of the Erasmus University Medical Center, Rotterdam, the Netherlands (MEC-2007-413).

The GINIplus and LISA studies were approved by the ethics committee of the Bavarian Board of Physicians (10 year follow up: 05100 for GINIplus and 07098 for LISA, 15 year follow up 10090 for GINIplus, 12067 for LISA).

The PANIC study protocol was approved by the Research Ethics Committee of the Hospital District of Northern Savo. All participating children and their parents gave informed written consent.

The YFS study protocol was approved by local ethics committees for contributing sites.

The RAINE study was approved by the University of Western Australia Human Research Ethics Committee.

### Phenotypes

Primary teeth exfoliate and are replaced by permanent teeth between 6 and 12 years of age. We aimed to separate caries status in primary and permanent teeth wherever possible using clinical information or age criteria, in line with our expectation that the genetic risk factors for dental caries might differ between primary and permanent dentition. For children in the mixed dentition we created two parallel case definitions, whilst in younger or older children a single case definition was sufficient.

All study samples included a mixture of children with dental caries and children who were caries-free, with varying degrees of within-mouth or within-tooth resolution. To facilitate comparison across these differing degrees of resolution all analysis compared children who were caries-free (unaffected) or had dental caries (affected). Missing teeth could represent exfoliation or delayed eruption rather than the endpoint of dental caries and therefore missing teeth were not included in classifying children as caries-free or caries affected.

In children aged 2.50 years to 5.99 years any individual with 1 or more decayed or filled tooth was classified as caries affected, with all remaining individuals classified as unaffected. In children aged 6.00 years to 11.99 years of age parallel definitions were determined for the primary dentition and permanent dentition respectively. Any individual with at least 1 decayed or filled primary tooth was classified as caries affected for primary teeth, while all remaining participants were classified as unaffected. In parallel, any individual with at least 1 decayed or filled permanent tooth was classified as caries affected for permanent teeth, while all remaining individuals were classified as unaffected. In children and adolescents aged 12.00 to 17.99 years of age any individual with 1 or more decayed or filled tooth or tooth surface (excluding third molar teeth) was classified as caries affected, with remaining individuals classified as unaffected.

Analysis was conducted in cross-section, meaning a single participant could only be represented in a single phenotype definition once. Where multiple sources of dental data were available for a single participant within a single phenotype definition window, the first source of data was selected (reflecting the youngest age at participation), in line with our expectation that caries status would be most heritable in the near-eruption period.

The sources of data used to create these phenotypic definitions are given in S3 Table. Within ALSPAC only, questionnaire responses were used to supplement data from clinical examination. The questions asked did not distinguish between primary and permanent teeth. Based on the age at questionnaire response we derived variables which prioritized responses from questionnaires before 6.00 years of age (thought to predominantly represent caries in primary teeth), and responses after 10.00 years of age (which might predominantly represent caries in permanent teeth). The final data sweep considered in this analysis targeted adolescents at age 17.50 years. Some participants responded to this after their eighteenth birthday. Data derived from this final questionnaire sweep were not included in the principal meta-analyses but were included in the GCTA heritability analysis.

### Genotypes and imputation

All participating studies used genetic data imputed to a comprehensive imputation panel. The 1000 genomes phase 1 version 3 panel (1KG phase 1 v3) was used as a common basis across 6 centers (GINIplus/LISA, GENR, GENEVA, YFS, PANIC, RAINE, (S1 Table). In ALSPAC, DNBC, COPSAC2000 and COPSAC 2010 the haplotype reference consortium (HRC v1.0 and v1.1) imputation panels were used. (S1 Table)

Each study performed routine quality control measures during genotyping, imputation and association testing (S2 Table). Further pre-meta-analysis quality control was performed centrally using the EasyQC R package and accompanying 1KG phase1 v3 reference data [34]. Minor allele count (MAC) was derived as the product of minor allele frequency and site-specific number of alleles (twice the site-specific sample size). Variants were dropped which had a per-file MAC of 6 or lower, a site-specific sample size of 30 or lower, or an impute INFO score of less than 0.4. Sites which reported effect and non-effect alleles other than those reported in 1KG phase 1 v3 reference data were dropped. Following meta-analysis, sites with a weighted minor allele frequency (MAF) of less than 0.5% were dropped, along with variants present in less than 50% of the total sample.

## Statistical analysis

### Association testing

Each cohort preformed GWA analysis using an additive genetic model. Caries status was modelled against genotype dosage whilst accounting for age at phenotypic assessment, age squared, sex and cryptic relatedness. Sex was accounted for by deriving phenotypic definitions and performing analysis separately within male and female participants, or by including sex as a covariate in association testing. Studies applied standard exclusions based on cryptic relatedness and ancestry, as described in S2 Table. In the GENR study association analysis included genetic principal components derived from the entire study population to account for ancestry in the multi-ethnic analysis, whilst the analysis of individuals of European ancestry included European population-specific genetic principal components [35]. The software and exact approach used by each study is shown in S2 Table.

### Meta-analysis

Results of GWA analysis within each study were combined in two principal meta-analyses, representing caries status in primary teeth and caries status in permanent teeth. For primary teeth, parallel meta-analyses were performed, one using results of multi-ethnic analysis in the GENR study and the other using results of European ancestry analysis in the GENR study. The GENR study did not have phenotypic data for permanent teeth, therefore the analysis of permanent teeth contained only individuals of European ancestry. Fixed-effects meta-analyses was performed using METAL [36], with genomic control of input summary statistics enabled and I test for heterogeneity. Meta-analysis was run in parallel in two centers and results compared. All available studies with genotype and phenotypic information were included in a one stage design, therefore there was no separate replication stage.

### Meta-analysis heritability estimates

For each principal meta-analysis population stratification and heritability were assessed using linkage disequilibrium score regression (LDSR) [37]. Reference linkage disequilibrium (LD) scores were taken from HapMap3 reference data accompanying the LDSR package.

### Within-sample heritability estimates

For comparison, heritability within the ALSPAC study was assessed using the GREML method [38], implemented in the GCTA software package [39], using participant level phenotype data and a genetic relatedness matrix estimated from common genetic variants (with effect allele frequency > 5.0%) present in HapMap3.

### Hypothesis free cross trait lookup

We used PLINK 2.0 [40] to clump meta-analysis summary statistics based on LD structure in reference data from the UK10K project. We then performed hypothesis-free cross-trait lookup of independently associated loci using the SNP lookup function in the MRBase catalog [41]. Proxies with an r^2^ of 0.8 or higher were included where the given variant was not present in an outcome of interest. We considered performing hypothesis free cross-trait genetic correlation analysis using bivariate LD score regression implemented in LDhub [42].

### Lookup in previously published pediatric caries GWAS

Previously published caries GWAS was performed within the GENEVA consortium, which is also represented in our meta-analysis. We therefore did not feel it would be informative to undertake lookup of associated variants in previously published results.

### Lookup in GWAS for adult caries traits

This analysis was planned and conducted in parallel with analysis of quantitative traits measuring lifetime caries exposure in adults (manuscript in draft).The principal trait studied in the adult analysis was an index of decayed, missing and filled tooth surfaces (DMFS). This index was calculated from results of clinical dental examination, excluding third molar teeth. The DMFS index was age-and-sex standardized within each participating adult study before GWAS analysis was undertaken. Study-specific results files were then combined in a fixed-effects meta-analysis [43]. In addition to DMFS, two secondary caries traits were studied in adults, namely number of teeth (a count of remaining natural teeth at time of study participation) and standardized DFS (derived as the number of decayed and filled surfaces divided by the number of natural tooth surfaces remaining at time of study participation). After age-and-sex standardization these secondary traits had markedly non-normal distribution and were therefore underwent rank-based inverse normal transformation before GWAS analysis and meta-analysis. We performed cross-trait lookup of lead associated variants in the pediatric caries meta-analysis against these three adult caries traits. As the unpublished analysis also contains samples which contributed to previously published GWAS, we did not feel it would be informative to undertake additional lookup in published data.

### Gene prioritization, gene set enrichment and association with gene transcription

Gene based testing of summary statistics was performed using MAGMA [44] with reference data for LD correction taken from the UK10K project and gene definitions based on a 50 kilobase window either side of canonical gene start:stop positions. Gene set enrichment analysis was considered using the software package DEPICT [45]. Tests for association between phenotype and predicted gene transcription were performed using MetaXcan [46] using a transcription prediction model trained in whole blood (obtained from the PedictDB data repository at http://predictdb.org/) [47]. Bonferroni correction was applied on the basis of approximately 7,000 independent gene based tests for 2 caries traits, giving an experiment wide significance level of approximately *p*<3.6e-06.

### Power calculations

Post-hoc power calculations were performed using the free, web-based tool Genetic Association Study (GAS) Power Calculator and the software utility Quanto (v1.2.4) (https://csg.sph.umich.edu/abecasis/gaspowercalculator/index.html, http://biostats.usc.edu/Quanto.html) [48]. Using the sample size and caries prevalence of the final meta-analysis samples, we calculated the minimum effect size required to have 80% discovery power at a significance level of 5.0e-08 for variants with MAF between 0.05 and 0.50. For primary teeth (17,037 individuals, 6,922 caries affected, prevalence 40.6%) we were able to detect variants with a minimal effect size (OR) between 1.13 and 1.37 for variants with MAF of 0.50 and 0.05, respectively (1.15 for MAF of 0.30) (S4 Fig, S5 Fig). For permanent teeth (13,353 individuals of which 5,875 were caries-affected, prevalence 44.0%) we had 80% power to detect variants with a minimal effect size (OR) between 1.15 and 1.43 for variants with MAF of 0.50 and 0.05, respectively (1.17 for MAF of 0.30). (S4 Fig, S5 Fig)

## Results

### Single variant results

Meta-analysis of caries in primary teeth in individuals of European ancestry included 17,037 individuals (6,922 affected) from 22 results files representing all 9 coordinating centers. After final quality control (QC), this meta-analysis included 8,640,819 variants, with mild deflation (genomic inflation factor (λ) = 0.994)(S1 Fig). Meta-analysis of caries in primary teeth which included individuals of multiple ethnicities in the GENR study included 19,003 individuals (7,530 affected) from 22 results files representing all 9 coordinating centers. There were 8,699,928 variants after final QC, with mild deflation in summary statustics (λ = 0.986)(S2 Fig). Analysis of caries status in permanent teeth included 13,353 individuals (5,875 affected) from 14 results files representing 7 coordinating centers. The sample size was smaller for permanent teeth as two coordinating centers did not have phentoype data for permanent teeth (RAINE and GENR), whilst the COPSAC group only had data for participants in the earlier birth cohort (COPSAC 2000). There were 8,734,121 variants afer final QC, with mild deflation in summary statistics (λ = 0.999)(S3 Fig).

The strongest evidence for association with caries in primary teeth was seen at rs1594318 (OR 0.85 for C allele, EAF 0.60, p = 4.13e-08) in the European ancestry meta-analysis (Figs 1,2 and 3, Table 1). This variant is intronic within *ALLC* on 2p25, a locus which has not previously been reported for dental caries traits. In the meta-analysis combining individuals of all ancestories this variant no longer reached genome-wide significance, although suggestive evidence persisted at rs1594318 (OR 0.868 for C allele EAF 0.60 p = 3.78e-07) and other intronic variants within *ALLC* in high linkage disequilibrium (Figure 3). For the permanent dentition the strongest statistical evidence for association was seen between caries status and rs7738851 (OR 1.28 for A allele, EAF 0.85, p = 1.63e-08). (Figs 1,2 and 4, Table 1). This variant is intronic within *NEDD9* on 6p24.

**Fig 1.**
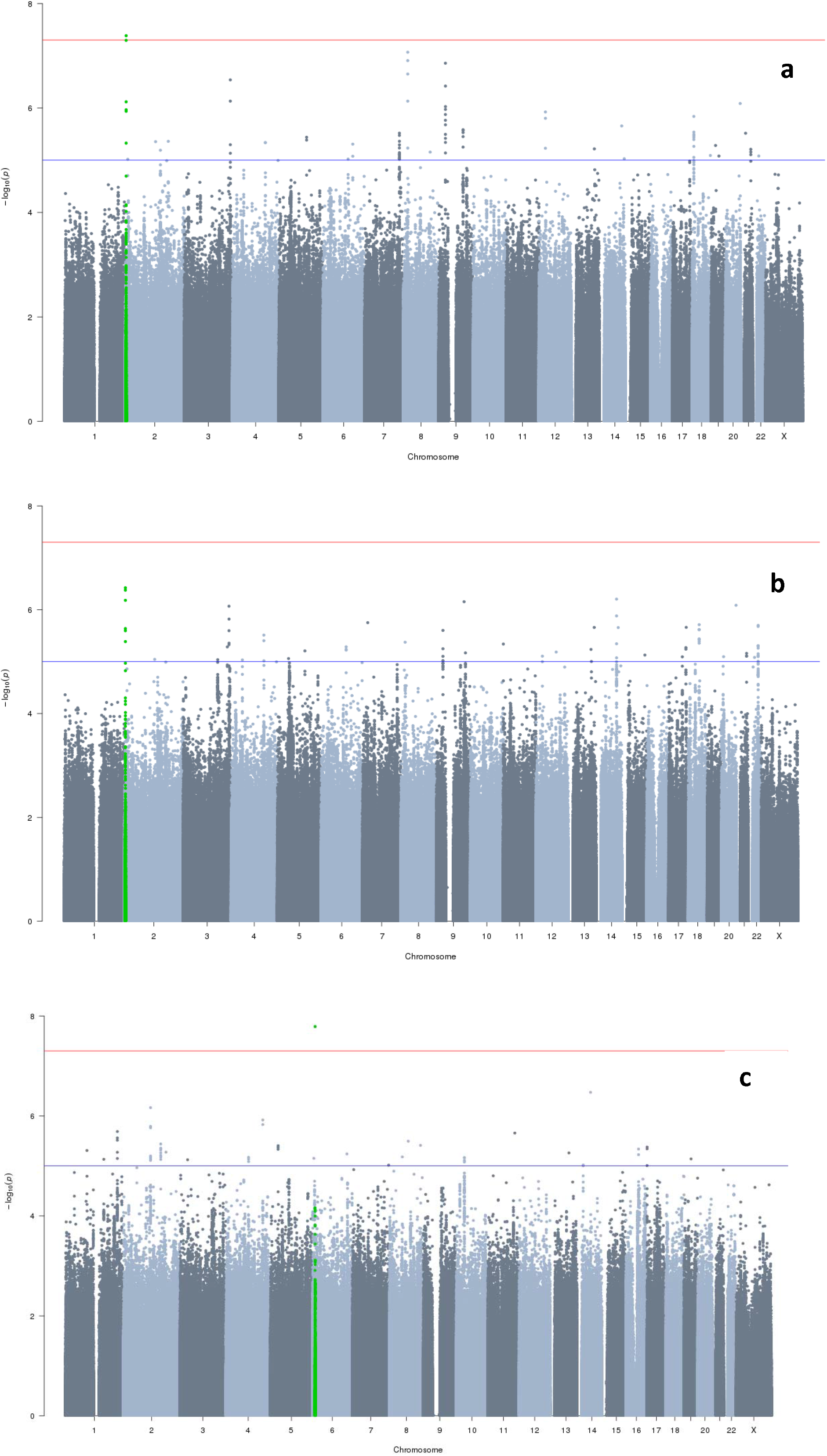
Manhattan plots for each principal meta-analysis. 1a: Caries in primary teeth (European ancestry), n samples = 17,036, n variants = 8,640,819, λ = 0.9944. Variants within 500Kb of rs1594318 are highlighted in green. 1b: Caries in primary teeth (multi-ethnic analysis), n samples = 19,003, n variants = 8,699,928, λ =0.9861. 2c: Caries in permanent teeth (European ancestry), n samples = 13,353, n variants = 8,734,121, λ =0.9991. Variants within 500Kb of rs7738851 are highlighted in green.

**Fig 2.**
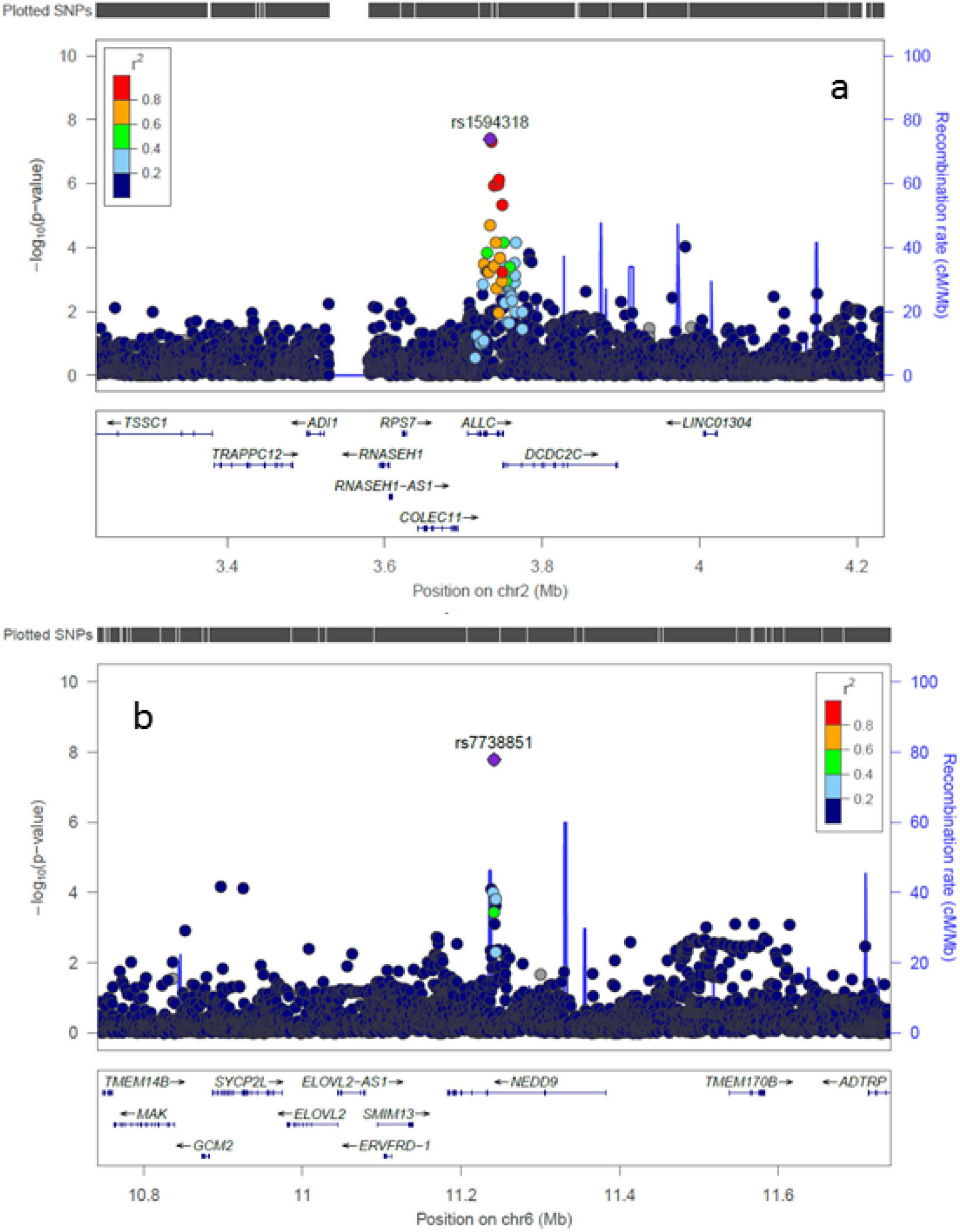
Regional association plots. 2a: Regional association plot for rs1594318 and caries in primary teeth (European ancestry meta-analysis. 2b: Regional association plot for rs7738851 and caries in permanent teeth.

**Fig 3.**
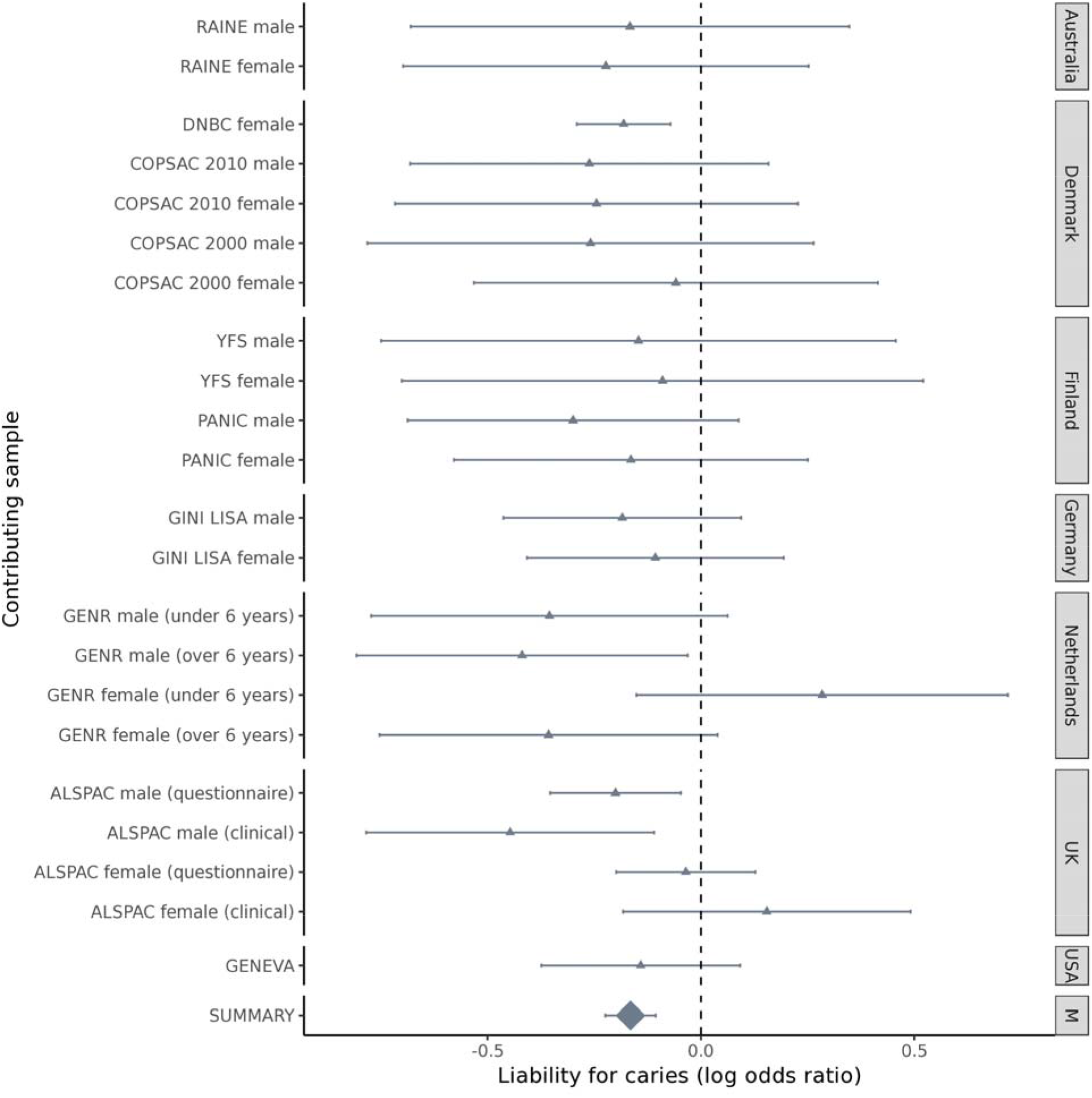
Forest plot for rs1594318 and caries in primary teeth. Effect sizes are expressed on a log odds ratio scale, grouped by geographical location. The summary estimate is from the fixed-effect metaanalysis of participants of European ancestry.

**Fig 4.**
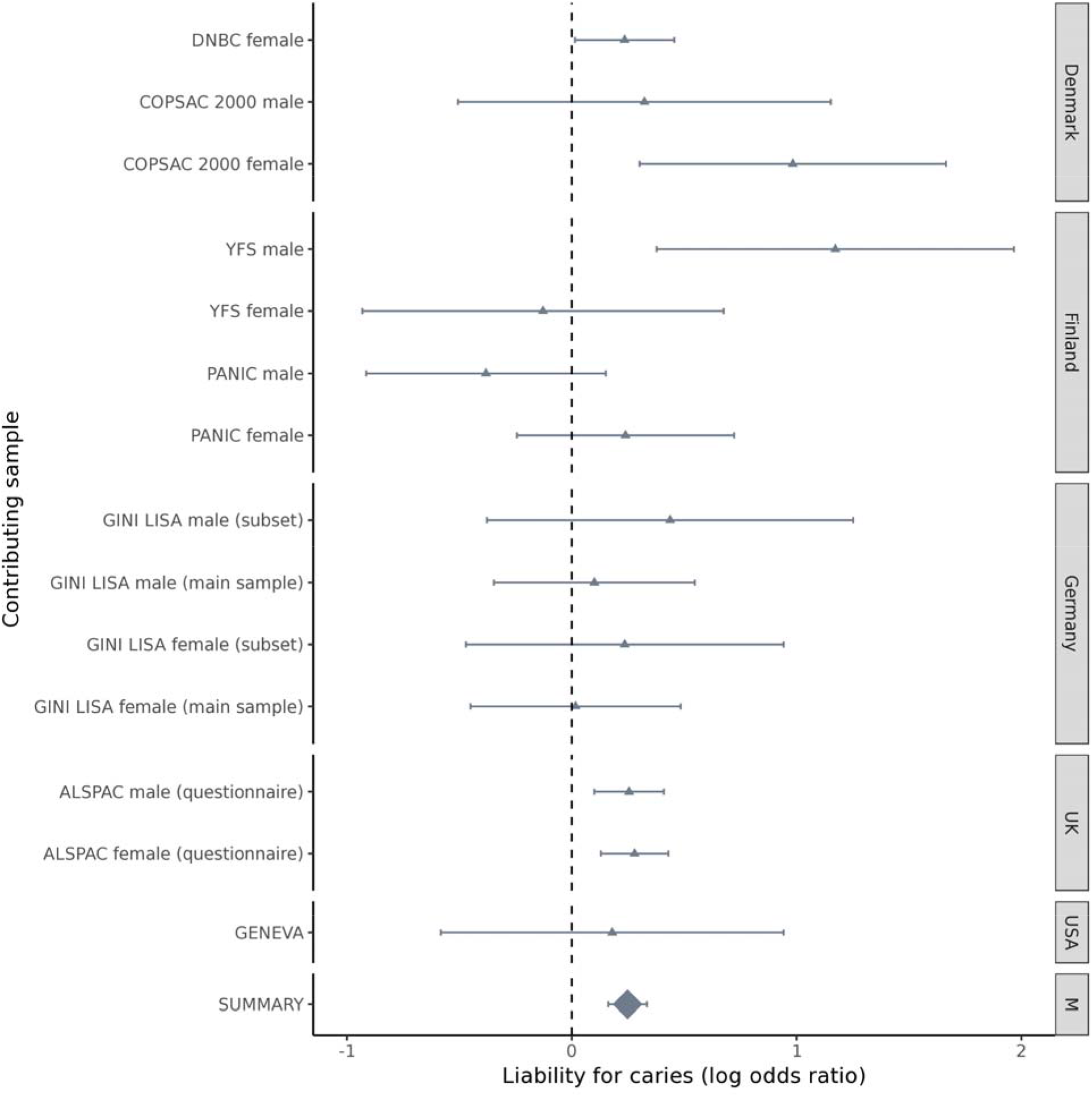
Forest plot for rs7738851 and caries in permanent teeth. Effect sizes are expressed on a log odds ratio scale, grouped by geographical location. The summary estimate is from fixed-effect metaanalysis.

**Table 1.**
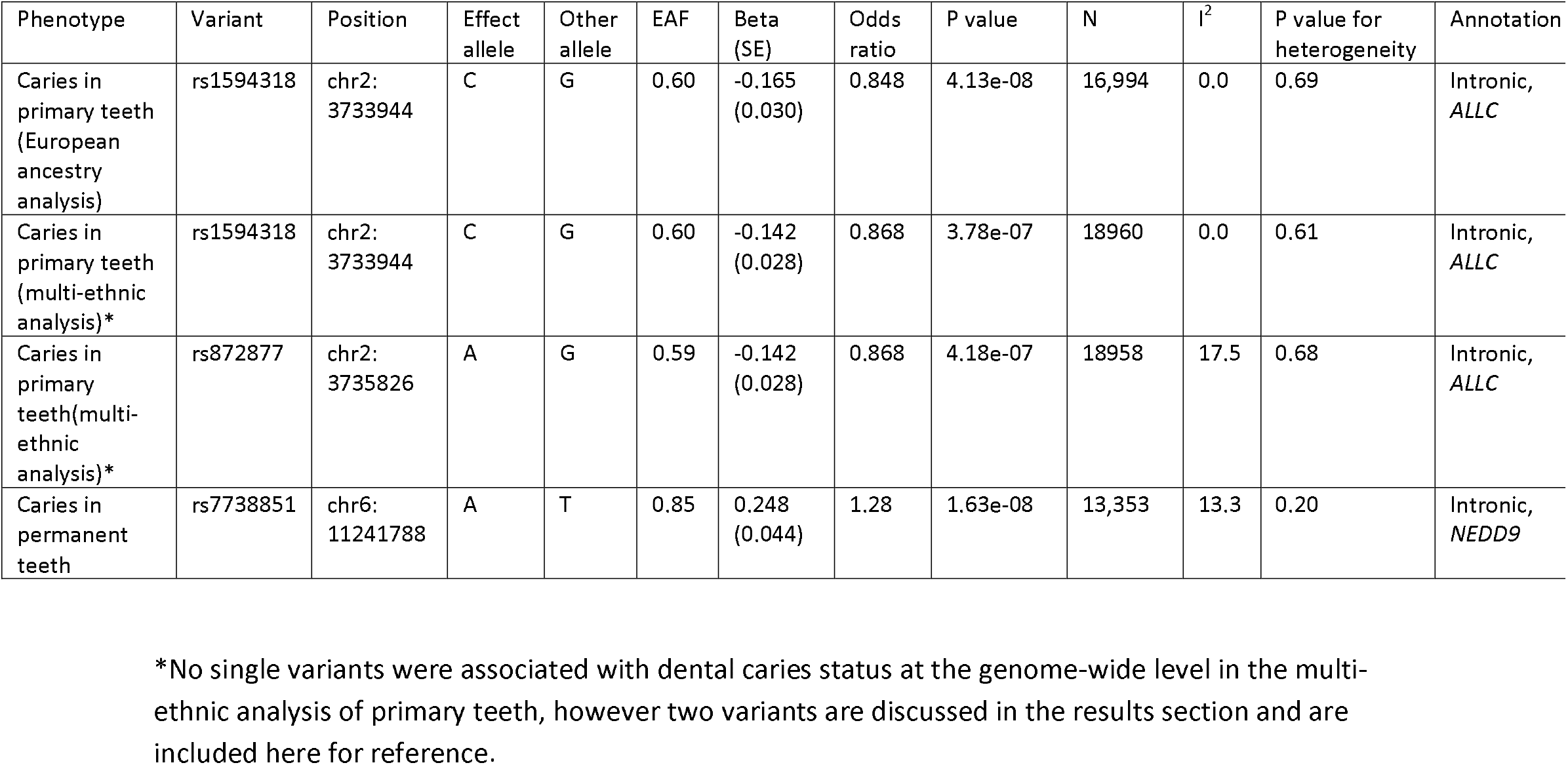
Lead associated single variants.

### Estimated heritabiltiy

Using participant level data in ALSPAC heritability was estimated at 0.28 (95% CI 0.09:0.48) and 0.17 (95% CI 0.02:0.31) for primary and permanent teeth respectively. Using summary statistics at the meta-analysis level produced point estimates near zero heritability, with wide confidence intervals (Table 2).

**Table 2.** Within-sample and meta-analysis heritability estimates

| Phenotype | Method | | Estimated $h^2$ (95% CI) | N |
| --- | --- | --- | --- | --- |
| Caries in primary teeth | GCTA GREML |  | 0.28 (0.09:0.48) | 7,230 |
|  | LDSR | All participants | 0.01 (0.00:0.06) | 19,003 |
|  |  | European ancestry only | 0.01 (0.00:0.07) | 17,036 |
| Caries in permanent teeth | GCTA GREML |  | 0.17 (0.02:0.31) | 6,657 |
|  | LDSR |  | 0.06 (0.00:0.12) | 13,353 |

### Cross-phenotype comparisons

Genome-wide mean chi squared was too low to undertake genome-wide genetic correlation using the LDSR method for caries in either primary or permanent teeth. Hypothesis-free phenome wide lookup for rs1594318 included 885 GWAS where either rs1594318 or a proxy with r > 0.8 was present. None of these traits showed evidence of association with rs1594318 at a Bonferroni-corrected alpha of 0.05. Lookup of rs7738851 and its proxies was performed against 662 traits, where similarly no traits reached a Bonferroni-corrected threshold. Hypothesis-driven lookup in adult caries traits revealed no strong evidence for persistent genetic effects into adulthood (Table 3).

**Table 3.** Lookup of lead associated variants

| Variant | Discovery trait | Risk increasing allele (discovery) | Cross trait lookup |  | P value | Effect per caries risk increasing allele (se) | N |
| --- | --- | --- | --- | --- | --- | --- | --- |
| rs1594318 | Caries in primary teeth (European ancestry meta-analysis) | G | Adult caries traits | DMFS (standard deviation of residuals of caries-affected surfaces) | 0.87 | -0.0015 (0.0092) | 26,790 |
|  |  |  |  | Number of teeth (inverse normal transformed residuals) | 0.60 | 0.0051(0.0098) | 27,947 |
|  |  |  |  | Standardized DFS (inverse normal transformed residuals) | 0.033 | -0.0195(0.0091) | 26,532 |
|  |  |  | Hypothesis free | (no traits meeting threshold for multiple testing) |  |  |  |
| rs7738851 | Caries in permanent teeth | A | Adult caries traits | DMFS (standard deviation of residuals of caries-affected surfaces) | 0.57 | -0.007 (0.011) | 26,791 |
|  |  |  |  | Number of teeth (inverse normal transformed residuals) | 0.63 | -0.0064 (0.013) | 27,949 |
|  |  |  |  | Standardized DFS (inverse normal transformed residuals) | 0.65 | -0.0054 (0.012) | 26,531 |
|  |  |  | Hypothesis free | (no traits meeting threshold for multiple testing) |  |  |  |

Adult caries traits were defined as follows. DMFS – a count of the number of decayed, missing or filled tooth surfaces. This count was residualized after regression on age and age squared and standard deviations of residuals calculated. Number of teeth – a count of the number of teeth in the mouth. This count was residualized after regression on age and age squared and residuals underwent inverse normal transformation. Standardized DFS. The number of decayed and filled surfaces was divided by the total number of tooth surfaces in the mouth. This ratio was residualized after regression on age and age squared and residuals underwent inverse normal transformation.

### Gene prioritization, gene set enrichment and association with gene transcription

Gene based tests identified association between caries status in the primary dentition and a region of 7q35 containing *TCAF1, OR2F2* and *OR2F1* (p=1.91e-06, 1.58e-06 and 1.29e-06, respectively). There were insufficient independently associated loci to perform gene set enrichment analysis using DEPICT for either of the principal meta-analyses. Association with gene transcription was tested but no genes met the threshold for association after accounting for multiple testing. The single greatest evidence for association was seen between increased transcription of *CDK5RAP3* and increased liability for permanent caries (*p*=3.94e-05). *CDK5RAP3* is known to interact with *PAK4* and *p14^ARF^*, with a potential role in oncogenesis [49, 50].

## Discussion

Dental caries in children and adolescents has not been studied to date using a large-scale, consortium based genome-wide meta-analysis aproach. Based on previous knowledge of the heritability of caries in young populations and from our understanding of other complex diseases, we anticipated that common genetic variants would be associated with dental caries risk with consistent effects across different cohorts. We found evidence for association between rs1594318 and caries in primary teeth. This variant showed weaker evidence for association in the multi-ethnic meta-analysis, potentially relating to different alelle frequencies across the different ethnic groups included in analysis. Frequency of the G allele is reported to vary between 0.24 in Asian populations to 0.42 in populations of European ancestry based on 1KGP allele fequencies. *ALLC* (Allanticase) codes the enzyme allantoicase, which is involved in purine metabolism and whose enzymatic activitiy is believed to have been lost during vertebrate evolution. Mouse studies suggest this loss of activity relates to low expression levels and low substrate affinity rather than total non-functionality [51]. Although there is some evidence that *ALLC* polymorphisms are associated with response to asthma treatment [52], there is limited understanding of the implications of variation in *ALLC* for human health, and it is possible that rs1594318 tags functionality elsewhere in the same locus.

For permanent teeth we found evidence for association between caries status and rs7738851, an intronic variant with *NEDD9* (neural precursor cell-expressed, developmentally down-regulard gene 9). *NEDD9* is reported to mediate integrin-initiated signal transduction pathways and is conserved from gnathostomes into mammals [53] [54]. *NEDD9* appears to play a number of functional roles in disease and normal development, including regulation of neuronal differentiation, development and migration [53, 55-59]. One such function involves regulation of neural crest cell migration [57]. Disruption of neural crest signalling is known to lead to enamel and dentin defects in animal models [60, 61] and might provide a mechanism for variation at rs7738851 to influence dental caries susceptibiltiy.

Traditionally risk assessment for dental caries in childhood has concentrated on dietary behaviours and other modifiable risk factors [62], with little focus on tooth quality. Although our understanding of the genetic risk factors for dental caries is incomplete, authors have noted that the evidence from previous genetic association studies tends to support a role for innate tooth structure and quality in risk of caries [63, 64]. If validated by future studies, the association with rs7738851 would provide further evidence for this argument, and may in the future enhance risk assessment in clinical practice.

The lookup of lead associated variants against adult caries traits provided no strong evidence for persistent association in adulthood. This might imply genetic effects which are specific to the near-eruption timepoint. An alternative explaination is that the variants identified in the present study represent false positive signals; although we see good consistency of effects across studies the statistical evidence presented is not irrefutable and there is no formal replication stage in our study.

The meta-analysis heritabiltiy estimates were lower than anticipated from either previous within-study heritability estimates [65] or the the new within-study heritabiltiy estimates obtained for this analysis. There are several possible explanations for this phenomenom. First, the methods used in the present analysis are SNP based which consistently underestimate heritability of complex traits relative to twin and family studies [66]. Second, meta-analysis heritabiliy represents the heritability of genetic effects which are consistent across populations. In the event of genuine differences in the genetic architecture of dental caries across strata of age, geography, environmental eposures or subtly different phenotypic meanings the meta-heritabiltiy estimate is not the same conceptually as the weighted average of heritabiltiy within each study.

More intuitively, genetic influences might be important within populations with relatively similar environments but not determine much of the overall diffrences in risk when comparing groups of people in markedly different environments. This view is consistent with existing literature from family based and candidate gene association studies suggesting the genetic architectre of dental caries is complex with multiple interactions. For example, gene-sex interactions are reported which change in magnitude between the primary and permanent dentition [67], genetic variants may have heterogeneous effects on the primary and permanent dentition [68] and environmental exposures such as fluoride may interact with genetic effects [69]. Finally, the aetiological relevance of specific microbiome groups appears to vary between different populations [70], suggesting genetic effects acting through the oral microbiome might also vary between populations. Unfortunately, this study lacks statistical power to perform meta-analyses stratified on these exposures, so does not resolve this particular question.

In line with any consortium based approach, the need to harmonize analysis across different collections led to some compromises.The phenotypic definitions used in this study do not contain information on disease extent or severity. Loss of information in creating these defintions may have contributed to the low statistical power of analysis. Our motivation for using simple definitions was based on the facts that a) case-control status simply represents a threshold level of an underlying continium of disease risk b) simple binary classifications facilitate comparison of studies with different assessment protocols and population risks and c) simple classifications have been used sucsessully in a range of complex phenotypes. Between participating centers there are differences in characteristics such as age at participation, phenotypic assesment and differences in the environment (such as nutrition, oral hygiene and the oral microbime) which might influence dental caries or its treatment, as reflected in the wide range of caries prevalence between different study centers. Potentially, this might lead to heterogenetiy in our meta-analyses. Although we see little evidence for heterogeneity in the top associated loci reported, it is possible that heterogeneity at other loci contributed to low study power and prevented more comprehensive single variant findings.

In the ALSPAC study we made extensive use of questionnaire derived data. This will systematically under-report true caries exposure compared to other studies as children or their parents are unlikely to be aware of untreated dental caries which would be evident to a trained assesor. We have explored some of these issues previously and shown that self-report measures at scale can be used to make meaningful inference about dental health in childhood [71]. We believe that misclassification and under-reporting in questionnaire data would tend to bias genetic effect estimates and heritability towards the null. Despite this we show evidence for heritabiltiy using these definitions and effect sizes at lead variants are comparable with effect sizes obtained using clinically assessed data (Figs 3,4).

As our power calculations showed, the sample size was sufficient to detect the identified variants associated at a genome wide significant level with caries in the primary teeth (rs1594318) and in permanent teeth (rs872877), where we observed relatively large effect sizes. For smaller effect sizes we were underpowered to identify association, and did not detect any variants with effect sizes (expressed as per-allele increased odds) smaller than 15% or 17% in the primary and permanent teeth, respectively. Caries is highly influenced by environmental factors and it is likely that individual genetic variants only have small effects on caries risk [3–6]. Much of the heritabiltiy of dental caries may be carried by variants with much smaller effects, as seen in other comparable complex traits [72]. Furthermore, some of the included studies had major differences in their caries prevalence, likely acting as a proxy for features affecting risk of caries. This may have introduced heterogeneity and reduced power to detect association, as discused further below.

One area of interest in the literature is the ability of genetics to guide personalized decisions on risk screening or identifying treatment modalities, and this is also true in dentistry. The genetic variants identified in this study are unlikely to be useful on their own in this context, given the modest effect sizes and low total heritability observed in our meta-analysis. We would suggest clinicians should continue to consider environment and aggregate genetic effects (for example, knowledege of disease patterns of close relatives) rather than specific genetic variants at this moment in time. Nevertheless, the findings of our study contribute to a better understanding of the genetic and bioloigcal mechanisms underlying caries suceptibility.

## Supporting information

**S1 Fig. Quartile-Quartile plot for caries in primary teeth (European ancestry analysis)**.

**S2 Fig. Quartile-Quartile plot for caries in primary teeth (multi-ethnic anlaysis)**.

**S3 Fig. Quartile-Quartile plot for caries in permanent teeth**.

**S4 Fig. Power calculations for caries in primary and permanent teeth**. This figure shows estimated power to detect association at a range of effect sizes for caries in the primary dentition (European ancestry analysis) and caries in the permanent dentition. Minor allele frequency is fixed at 0.40 and significance level is fixed at 5e-08.

**S5 Fig. Power calculations at a range of minor allele frequencies**. This figure shows estimated power to detect association at a range of minor allele frequencies and effect sizes for caries in the primary dentition (European ancestry analysis) (a) and caries in the permanent dentition (b). Significance level is fixed at 5e-08.

**S1 Table. Participating studies**. This table describes the participating cohorts and analytical centers

**S2 Table. Genotyping and quality control**. This table describes genotyping, quality control, imputation and association testing protocals follwod at each participating center.

**S2 Table. Phenotype summaries**. This table describes the prevalance of caries and demographics of participants within each stratified results file.

## Acknowledgements

We are very grateful to the children and families who agreed to participate in the contributing studies, without whom this research would not be possible. We would like to acknowledge the role of Mark McCarthy and the Early Growth Genetics consortium in recruiting studies which contributed to this analysis.

For ALSPAC, we are extremely grateful to all the families who took part in this study, the midwives for their help in recruiting them, and the whole ALSPAC team, which includes interviewers, computer and laboratory technicians, clerical workers, research scientists, volunteers, managers, receptionists and nurses.

The authors are grateful to the Raine Study participants and their families, and to the Raine Study research staff for cohort coordination and data collection. The authors gratefully acknowledge the assistance of the Western Australian DNA Bank (National Health and Medical Reserach Council of Australia National Enabling Facility). We would also like to acknowledge the Raine Study participants for their ongoing participation in the study, and the Raine Study Team for study co-ordination and data collection.

